# CoMPHI: A Novel Composite Machine Learning Approach Utilizing Multiple Feature Representation to Predict Hosts of Bacteriophages

**DOI:** 10.1101/2024.07.29.604684

**Authors:** Shreyashi Bodaka, Onkar Malgonde

## Abstract

Phage therapy has reemerged as a compelling alternative to antibiotics in treating bacterial infections, especially for superbugs that have developed antibiotic resistance. The challenge in the broader application of phage therapy is identifying host targets for the vast array of uncharacterized phages obtained through next-generation sequencing. To solve this issue, this paper introduces an innovative Composite Model for Phage Host Interaction, CoMPHI, to predict phage-host interactions by combining the accuracy of alignment-based methods with the efficiency and flexibility of machine learning techniques. The model initially generates multiple feature encodings from nucleotide and protein sequences of both phages and hosts to enhance prediction accuracies. It is further enriched by incorporating alignment scores between phage-phage, phage-host, and host-host, creating a composite model. During the 5-fold cross-validation, the composite model exhibited an Area Under the ROC Curve (AUC) of 94%, 96.4%, 96.5%, 96.6%, 96.6%, and 96.7% and accuracy of 92.3%, 93.3%, 93.6%, 94%, 94.9%, and 95.1% at the Species, Genus, Family, Order, Class, and Phylum levels, respectively. A comparative analysis revealed a 6-8% increase in model performance due to the inclusion of alignment scores. Additionally, an ablation study highlighted that including both nucleotide and protein sequences from both phages and hosts increased the prediction accuracy of the model. Another ablation study provided evidence that phage-host and host-host alignment scores, combined with phage-phage scores, equally contributed to enhancing the composite model’s performance. In conclusion, this paper presents a robust and comprehensive composite model advancing the use of phage therapy in modern medicine.

## Introduction

Antimicrobial resistance (AMR) was declared one of the top 10 global health threats by the World Health Organization (WHO). Antibiotics, considered a cornerstone of modern healthcare, are under threat from antibiotic resistance, which has emerged as a significant global public health and socioeconomic issue. An estimated 4.95 million deaths were attributed to bacterial antibiotic resistance, with 1.27 million deaths being specifically linked to bacterial AMR in 2019 (Murray et al., 2022). The World Bank projected that up to 3.8% of the global gross domestic product could be lost due to AMR by 2050 (Jonas et al., 2017). Among drug-resistant microbes, a significant threat is posed by the group known as ESKAPEE, an acronym for Enterococcus faecium, Staphylococcus aureus, Klebsiella pneumoniae, Acinetobacter baumannii, Pseudomonas aeruginosa, Enterobacter spp., and Escherichia coli. These pathogens comprise high to critically drug-resistant strains and fall into the WHO’s Critical Priority I and II categories. The pharmaceutical industry currently regards antibiotic development as financially imprudent (Safir et al., 2020) due to economic and regulatory barriers, resulting in diminished interest in this critical area (Bartlett et al., 2013). This circumstance heightens the imminent threat of entering a post-antibiotic era (Bartlett et al., 2013).

Phage therapy emerges as a compelling alternative to antibiotics in treating bacterial infections, particularly in combating superbugs that have developed resistance to traditional antibiotics. (Saw & Song, 2019). Phages are highly specific to the type of host they can infect (Saw & Song, 2019). This specificity implies that a particular phage would only target a particular strain or species of the host. Currently culture-based or in vitro methods are employed to characterize and isolate phages that lyse their specific hosts. However, this method is resource-, labor-, and time-intensive, heavily dependent on the lytic cycle, limited to hosts that can be cultivated, not suited to be applied in large-scale or complex environments, and has low efficiency (Hyman, 2019). Leveraging state-of-the-art metagenomic sequencing and advanced bioinformatics, in silico prediction of putative hosts for metagenomic sequenced phages can accelerate and broaden the application of phage therapy in modern medicine. Such predictions are based on genomic signals arising from the coevolution and/or arms race between phages and hosts. These can be broadly categorized as alignment-based and alignment-free/machine learning methods.

### Alignment-Based Methods

Alignment-based methods leverage genomic and proteomic sequence homology/similarity to predict the host range of a phage (Versoza & Pfeifer, 2022). Host genomes undergo modifications during coevolution and/or arms race with phages, involving integration of prophages, auxiliary metabolic genes (AMGs), shared tRNAs, and clustered regularly interspaced short palindromic repeats (CRISPRs) with CRISPR-associated protein (Cas) spacers (Barrangou et al., 2007; Edwards et al., 2017; Hampton et al., 2020, Coclet & Roux, 2021). These shared sequence similarities serve as indicators of either successful infection or evasion from host defenses. Widely utilized tools for predicting phage-host interaction through sequence similarity and homology searches include Blastn and Blastx (Altschul et al., 1990). While this method can achieve high accuracy by controlling the alignment threshold, its recall is limited due to the constraints of sequences in the search database (Ahlgren et al., 2017). Additionally, it struggles to adapt to evolutionary shifts in phage and host genomes (Hall et al., 2013).

### Alignment-Free/Machine Learning Methods

Alignment-free methods operate by extracting patterns and compositions from labeled empirical training data, employing statistical and/or probabilistic techniques, and do not rely on the alignment of sequences. The similarity in composition and/or patterns between phage and host is a result of phage genome adaptations to manipulate host translation machinery or to evade host restriction-modification mechanisms (Roux et al., 2015). Machine learning and deep learning methods utilize these pattern and composition features to predict phage-host interaction. The efficacy of a machine learning model hinges on capturing diverse genomic signals arising from intricate interactions between phages and hosts within a feature set (Li & Zhang, 2022).

While numerous studies have explored various aspects of phage-host genomic signals as feature sets for machine learning models, the focus has primarily been on either phage nucleotide sequences or specific proteins, such as WIsH (Galiez, 2017). Remarkably, only a limited number of studies have integrated both. Additionally, very few studies have delved into feature sets derived from both phages and hosts, overlooking the evident close co-evolution of these entities. Furthermore, encoding genetic sequences using a specific method may highlight specific composition properties or patterns associated with their function, potentially overlooking functionalities due to data sparsity. Consequently, machine learning algorithms might miss features related to other functionalities within the sequences. This gap underscores the potential for enhanced model accuracy through a more comprehensive exploration of combined phage and host genomic features. Within these broader categories, alignment-based methods demonstrate high accuracy but low recall whereas alignment-free methods exhibit higher recall, but lower precision compared to alignment-based methods. *This paper introduces a novel composite model for predicting phage-host interactions, hypothesizing that capitalizing on the accuracy of alignment-based methods and the recall and flexibility of machine-learning techniques will improve its performance further than the current literature.* The model first utilizes multiple feature encodings from both nucleotide and protein sequences of phages and hosts. Then it leverages similarity scores from alignment-based methods for phage-phage, phage-host, and host-host interactions, along with the machine learning algorithm to predict the interaction probabilities between phages and hosts.

## Materials and Methods

### Dataset Collection and Pre-Processing

The dataset contained genomes of phages and hosts and phage-host interaction downloaded from the National Center of Biotechnology Information (NCBI) RefSeq bacteriophage database available as of August 2023 (US National Library of Medicine). This included 3,629 unique phage-host interactions between 3,629 phages and 815 hosts. Only phages that infect bacteria along with their hosts were selected and incomplete genomes of nucleotides and protein sequences were removed. To reduce bias due to the over-representation of a particular phage, data redundancy was removed by clustering phage genomes using CD-HIT (Fu et al., 2012) at a 95% identity match resulting in a dataset containing 3,018 unique phage-host interactions between 3,018 phages and 353 hosts. Then phage-host interactions with incomplete host taxonomy were removed from the data resulting in a final data set with 2,948 unique phage-host interactions between 2,948 phages and 256 hosts. As there is no laboratory-tested negative phage-host interaction data, negative interaction data are generated using phage-host interactions that are not included in this final cleansed dataset. The NCBI datasets tool was utilized to collect taxonomy data. A custom Python script was used to mass retrieve taxonomy data from phylum to genus by inputting each host name into the tool at https://api.ncbi.nlm.nih.gov/datasets/v2alpha/taxonomy, which returned an Excel file with the taxonomy data.

### Composite Model Training, Validation and Testing

The composite model comprises primarily three key components:

**1. Generation of Alignment Bit Scores**: This involves creating alignment bit scores for phage-phage, host-host, and phage-host interactions.
**2. Generation of Multiple Feature Encodings**: This step focuses on generating multiple feature encodings for the nucleotides and proteins of both phages and hosts.
**3. Construction of a Composite Machine Learning Model**: In this stage, a composite machine learning model is developed by combining alignment-free prediction using machine learning with the alignment-based bit scores.

### Generation of Alignment Bit Scores

Due to the process of co-evolution, phages and their hosts share common genetic elements. Consequently, there is a strong probability that a closely related host will be susceptible to the same phage, or that similar phages will target the same host. To leverage this close association, alignment bit scores were acquired for phage-host, host-host, and phage-phage interactions using NCBI BLAST (Altschul et al., 1990). Phage–phage alignment bit scores are acquired by establishing a reference phage database that includes all phages in the dataset. Each phage nucleotide in the dataset undergoes a search against this reference phage database using BLASTn with an e-value of 0.0001. The bit score of the maximum hit in this search (after excluding matches to itself) is documented as the phage–phage alignment score for the respective phage forming an array BIT_PP of dimension N, where N is the total number of unique phages in the dataset. Similarly, phage–host alignment bit scores are obtained by establishing a reference host database encompassing all hosts in the dataset. Each phage nucleotide in the dataset is then queried against this host reference database using BLASTn with an e-value of 0.0001. The bit score of the maximum hit in this search is recorded as the phage– host alignment score for the corresponding phage, resulting in an array BIT_PH of dimension NxM, where N is the total number of unique phages, and M is the total number of unique hosts in the dataset. Lastly, a search involving all hosts in the dataset against the reference host database, with an e-value of 0.0001, is conducted to register the host–host alignment scores. This process forms an array BIT_HH of dimension M, where M is the total number of unique hosts in the dataset.

### Generation of Multiple Feature Encodings

Utilizing multiple representations and a wider array of features extracted from nucleotides and proteins of both the phage and hosts enables harnessing complementary genetic signals from various levels of molecular interaction contributing to an enhanced accuracy in phage-host prediction. To encode nucleotide sequences, feature encodings that are agnostic to the length of nucleotides were used to mitigate the influence of sequence length on bias. Following a similar approach as Li et al. (2021), feature encodings for protein sequences were generated using the iLearn tool (Chen et al., 2020). The list of these encodings can be found in Table 1.

**Table 1.**
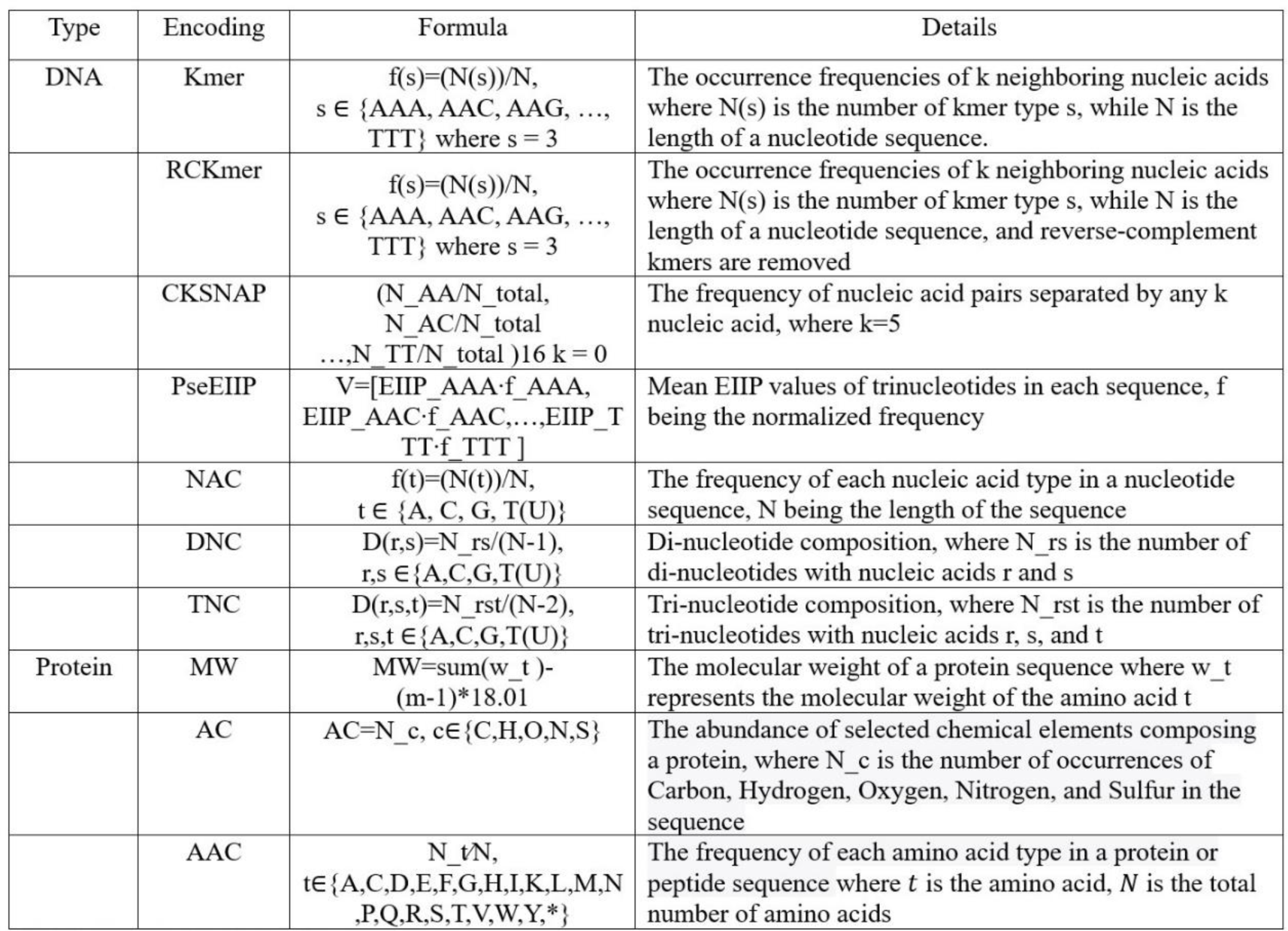
Details of Nucleotide and Protein Features.

As each phage or host consists of multiple protein sequences, six operators (mean, median, standard deviation, variance, maximum, and minimum) were employed to aggregate features derived from these multiple protein sequences. All feature encodings were normalized employing the min-max data normalization method, ensuring that the feature values fall within the range of 0 to 1. The feature encodings were consolidated into two feature vectors for each of the phage/host: one for nucleotide sequences and another for proteins.

To test deep learning models, the features were transformed into images. The individual sequential feature vectors originating from both phages and hosts underwent initial normalization employing the min-max data normalization method, ensuring that the feature values fall within the range of 0 to 1. These normalized vectors were then reshaped into an n x n array placing values into the array where n satisfies the condition: (n−1) ×(n−1) <N and N≤n×n. In cases where N≤n×n, padding is applied by introducing zeros to the remaining n×n−N entries (Xu et al., 2020). A bilayer architectural framework, incorporating nucleotide and protein layers, was devised by stacking feature vectors derived from phages and hosts.

### Construction of a Composite Machine Learning Model

Considering the exponential growth in genomic data and the objective of utilizing a single model for all viral genomes, a comparative study between several algorithms was conducted. Identifying key candidate algorithms was based on a review of the literature. Each algorithm differs based on underlying principles, assumptions, and approaches. These algorithms are also easy to implement and readily available via machine learning packages across multiple platforms/programming environments. Machine learning models that were evaluated are Logistic Regression (LR), Support Vector Machine (SVM), Neural Networks (NN), Decisions Tree (DT), K-Nearest Neighbor (KNN), Random Forest (RF), and Convolution Neural Network (CNN). After evaluating the model performance, considering the principles of parsimony and computational complexity, RF was identified as the superior algorithm for predicting the putative host of a phage as shown in Figure 2. This analysis utilized feature vectors extracted from the nucleotide and protein sequences of both phages and hosts. Probabilistic classifier RFs were constructed with 100 trees, considering 1,004 features for possible splitting at each node. Next, variable importance was estimated using the impurity method to access the contribution percentages of each of the features.

To further enhance each of the evaluated machine learning model’s performance, alignment-based bit scores between phage-phage, phage-host, and host-host were integrated with the machine learning prediction probabilities using a weighted sum as shown in equation 1. The host with the highest probability score after integrating the alignment bit scores was selected as the putative host of the phage.

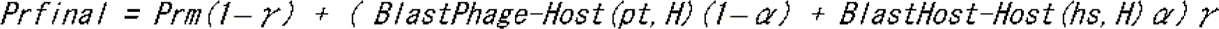

*Prm* is the prediction probabilities *Prm*=[*Prifori*=1,…*M*] from the RF model computed for all hosts, with *M* representing the total number of hosts in the dataset, and *Pri* denoting the prediction probability for the *i-th* host.

- *H* is all hosts in the dataset.
- *pt* represents the input/testing phage.
- *hs* represents the host of the most similar phage in the dataset based on BIT_PP
- *BlastPhage-Host* is using the BIT_PH scores between the phage and host (Figure 1E)
- *BlastHost-Host* is BIT_HH scores where the comparing host is the host of the top match from BIT_PP (Figures 1B, 1C, 1D)

**Figure 1.**
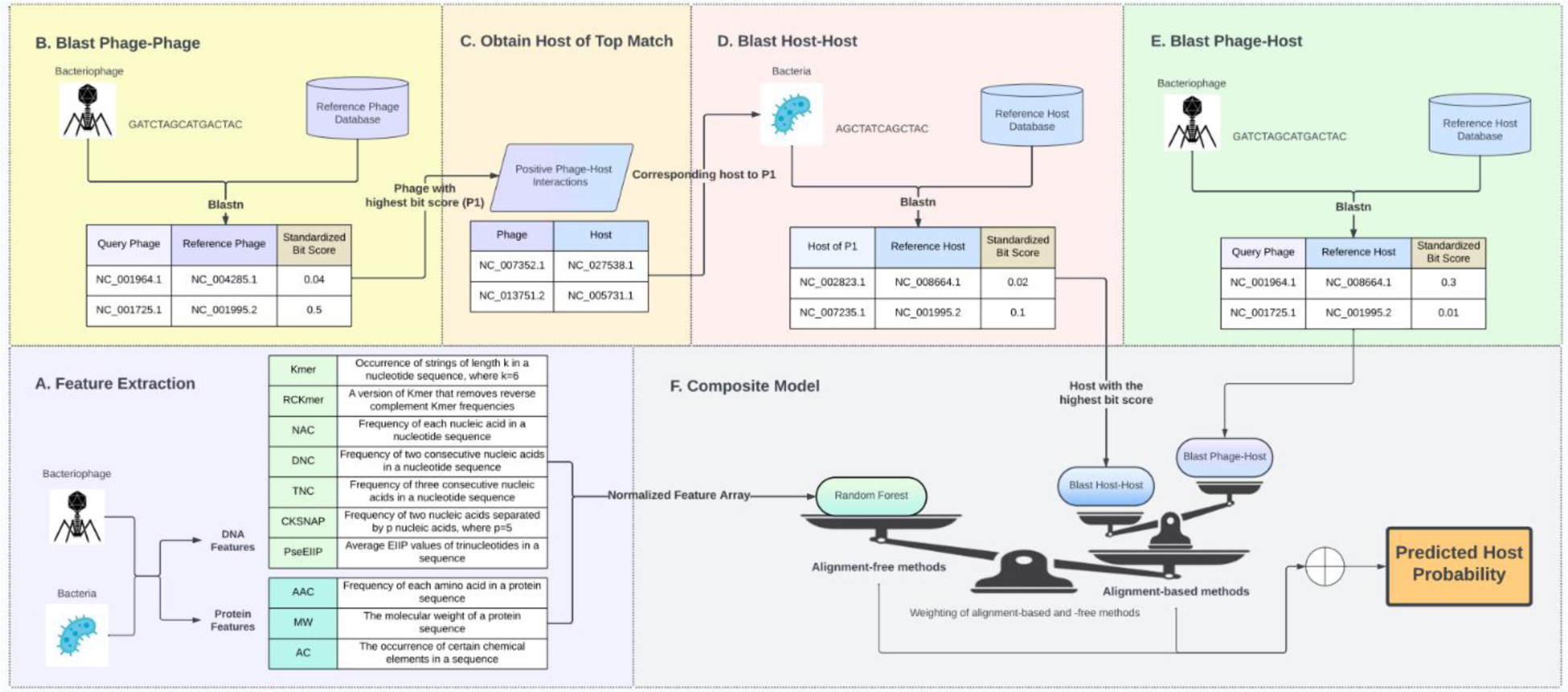
Composite Model Design. Figure 1A: Nucleotide and protein features from phage and host. Figure 1B, 1C, and 1D: Alignment score matrices BIT_PP, BIT_HH, and BIT_PH. Figure 1F: Composite Model.

### Model Optimization

Grid search was employed to fine-tune hyperparameters for the RF model. The optimal hyperparameter configuration identified through the grid search consisted of using 100 trees with a maximum depth of 20, a minimum number of samples required to split a node-set to 10, a minimum number of samples required at a leaf node set to 4, the maximum features set as sqrt, and a random state of 42.

Another iteration of the grid search was executed at 0.1 increments to determine the optimal weights for the contributions of machine learning predictions and alignment bit scores in the model. From this grid search, alpha and gamma values of 0.9 and 0.4, respectively, were identified.

## Results

### Model validation and performance

#### Performance of machine learning models

To measure performance, the following measures were used: accuracy (Acc), sensitivity (Sen), specificity (Spe), and area under the receiver-operating characteristic curve (AUC). 5-fold cross-validation was used on the entire dataset to ensure the generalization of the model and to assess model performance. Furthermore, to test the model on unseen data, testing was performed with the entire dataset using a randomized 70-30 split. This testing was repeated 10 times, and the averages were calculated to take care of data bias. Among all the algorithms included in the comparative analysis, RF and CNN demonstrated the best performance, at an accuracy and AUC of 86.5%, and 88.5% respectively for RF and 83.6%, and 88% respectively for CNN as illustrated in Figure 2. However, RF is a better algorithm due to its interpretability, computational simplicity, automatic feature importance, and simpler pre-processing of features. These metrics were within 2-3% of the 5-fold cross-validation further proving the robustness of the RF model on unseen data.

**Figure 2.**
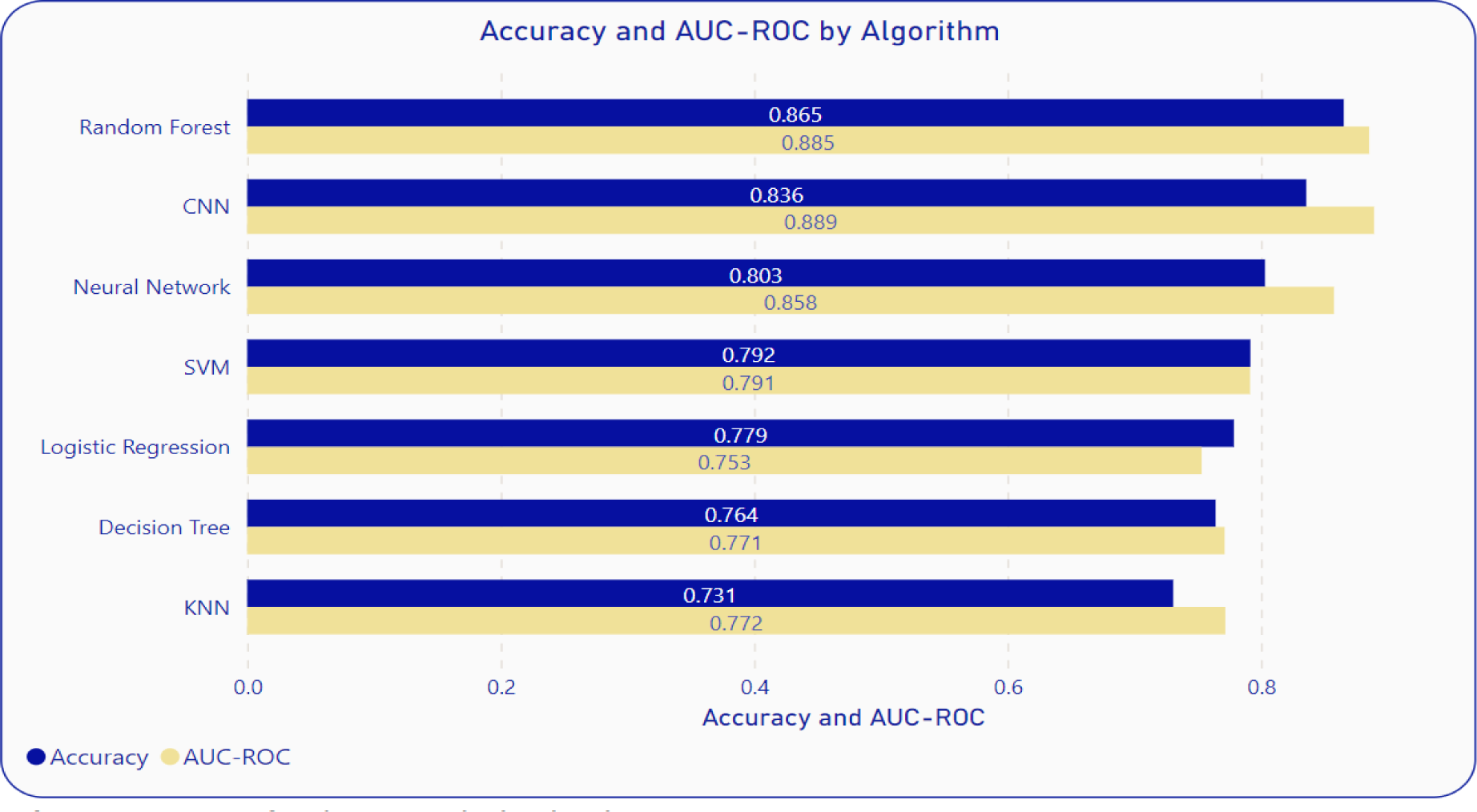
Performance Comparisons of RF with Other Algorithms

#### Performance of Composite Models

The machine learning model’s performance is improved by incorporating alignment scores. A random 70-30 split of the entire dataset was employed to assess the composite model. This testing process was iterated 10 times, and the averages were computed to address potential data bias. This resulted in AUC of 94%, 96.4%, 96.5%, 96.6%, 96.6%, and 96.7% and accuracy of 92.3%, 93.3%, 93.6%, 94%, 94.9%, and 95.1% at the Species, Genus, Family, Order, Class, and Phylum levels, respectively, for the RF composite model (as shown in Figure 3). When compared to the model utilizing only the machine learning algorithm, the composite model demonstrates approximately a 6-point higher performance, as illustrated in Figure 3. 5-fold cross validation using the entire dataset on the composite model also led to accuracies in 2-3% of 70-30 testing. The sensitivity and specificity were also improved by adding alignment-based scores (Figure 3), showing that alignment-based scores aid in improving True Positive Rate and True Negative Rate.

**Figure 3.**
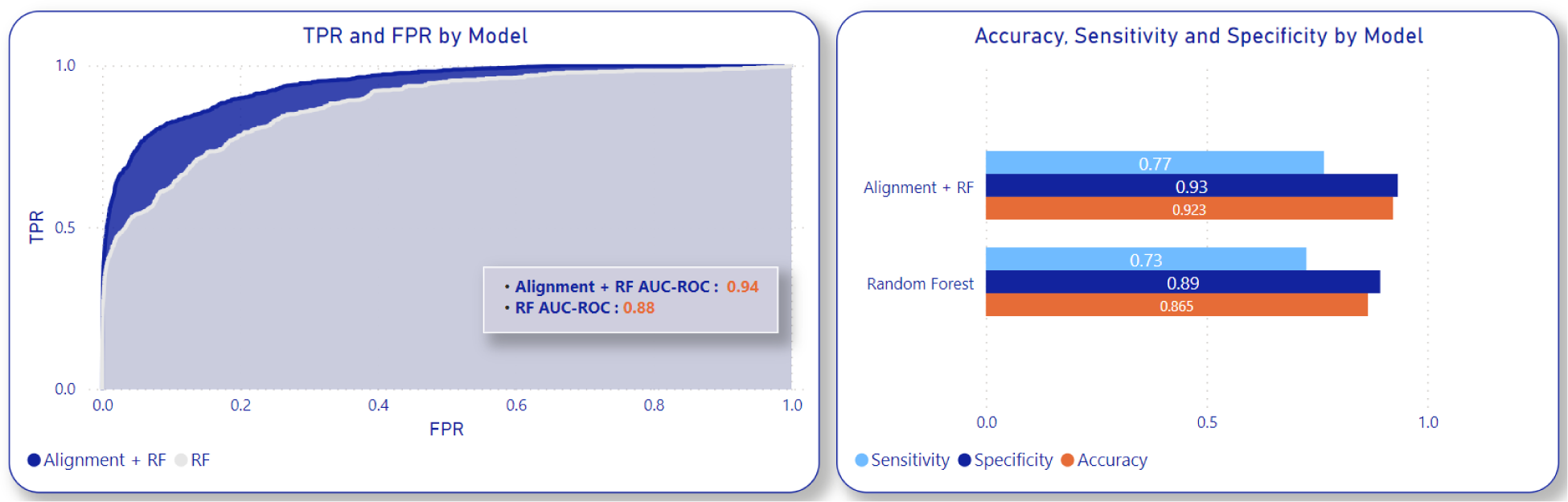
Composite Model Performance Compared to RF Alone

### Ablation Study

To understand the contributions of each of the components in the composite model with multiple features, an ablation analysis was conducted to compare the impact of including or excluding features. The models were evaluated based on the following feature combinations: only nucleotide features from phages, only protein features from phages, only nucleotide features from hosts, only nucleotide features from both phages and hosts, only protein features from both phages and hosts, and a combination of nucleotide and protein features from both phages and hosts. This analysis led to the conclusion that utilizing both nucleotide and protein features from both phages and hosts resulted in the highest prediction accuracies, as depicted in Figure 4. Furthermore, nucleotide features had a greater influence on the prediction than proteins. Host nucleotides and/or proteins contributed more to the model performance than the phage nucleotides and/or proteins. Another ablation analysis was conducted to assess the impact of individual alignment scores—phage-phage, phage-host, and host-host—on the composite model. In this analysis, the following combinations were tested using 70-30 randomized data split on the entire dataset. The test was repeated 10 times, and scores were averaged across these tests: BlastPhage-Host with the RF model, BlastHost-Host with the RF model, and BlastHost-Host with BlastPhage-Host and the RF model. As shown in Figure 5, the composite model using BlastHost-Host with BlastPhage-Host and the RF model scored the highest, indicating that utilizing all alignment scores helps increase accuracy.

**Figure 4.**
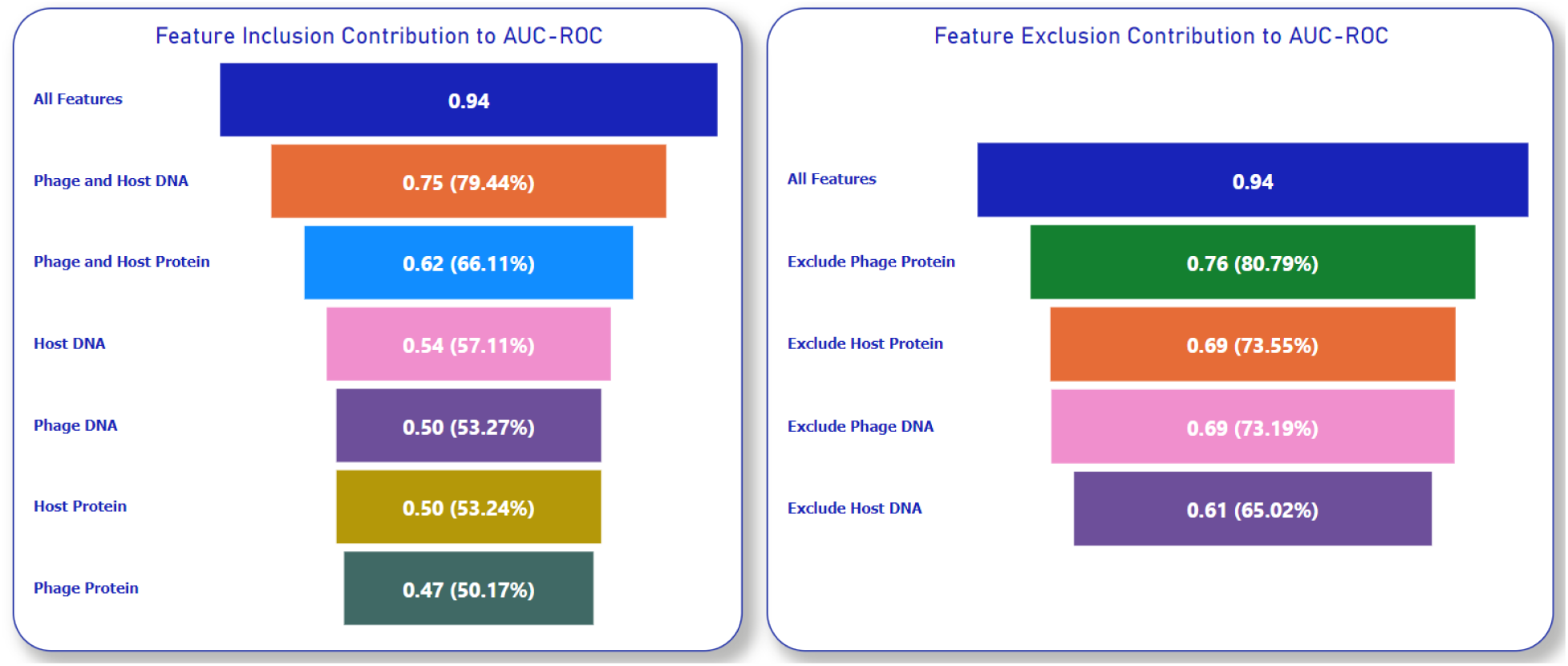
Composite Model Performance with Inclusion and Exclusion of Nucleotide and Protein Features

**Figure 5.**
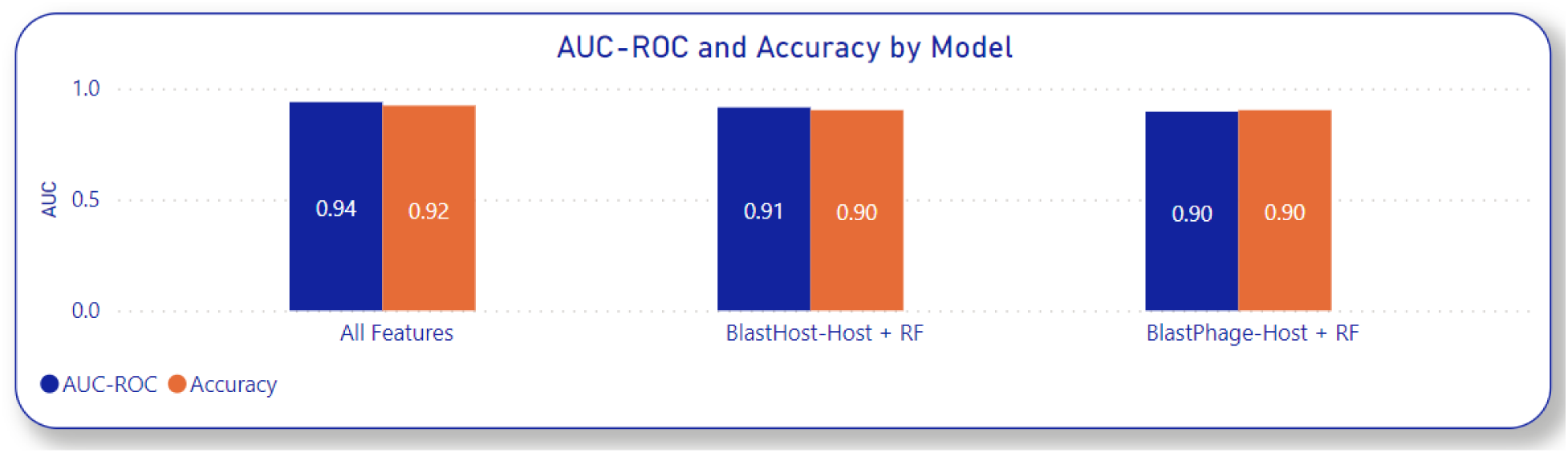
Composite Model Performance with Inclusion of Alignment Score Components

## Discussion

As proposed in this paper and proven by the ablation studies, composite features from both alignment-based and alignment-free methods as well as including multiple feature encodings from both nucleotides and proteins of phage and host have significantly improved the model’s performance in predicting phage-host interactions. To further gain a deeper understanding of the contribution of each alignment-free feature encoding, variable importance was assessed using the impurity method in the RF model. This estimation further proved the results of the ablation study that the nucleotide features made a greater contribution to the prediction accuracy compared to the protein features as shown in Figure 6.

**Figure 6.**
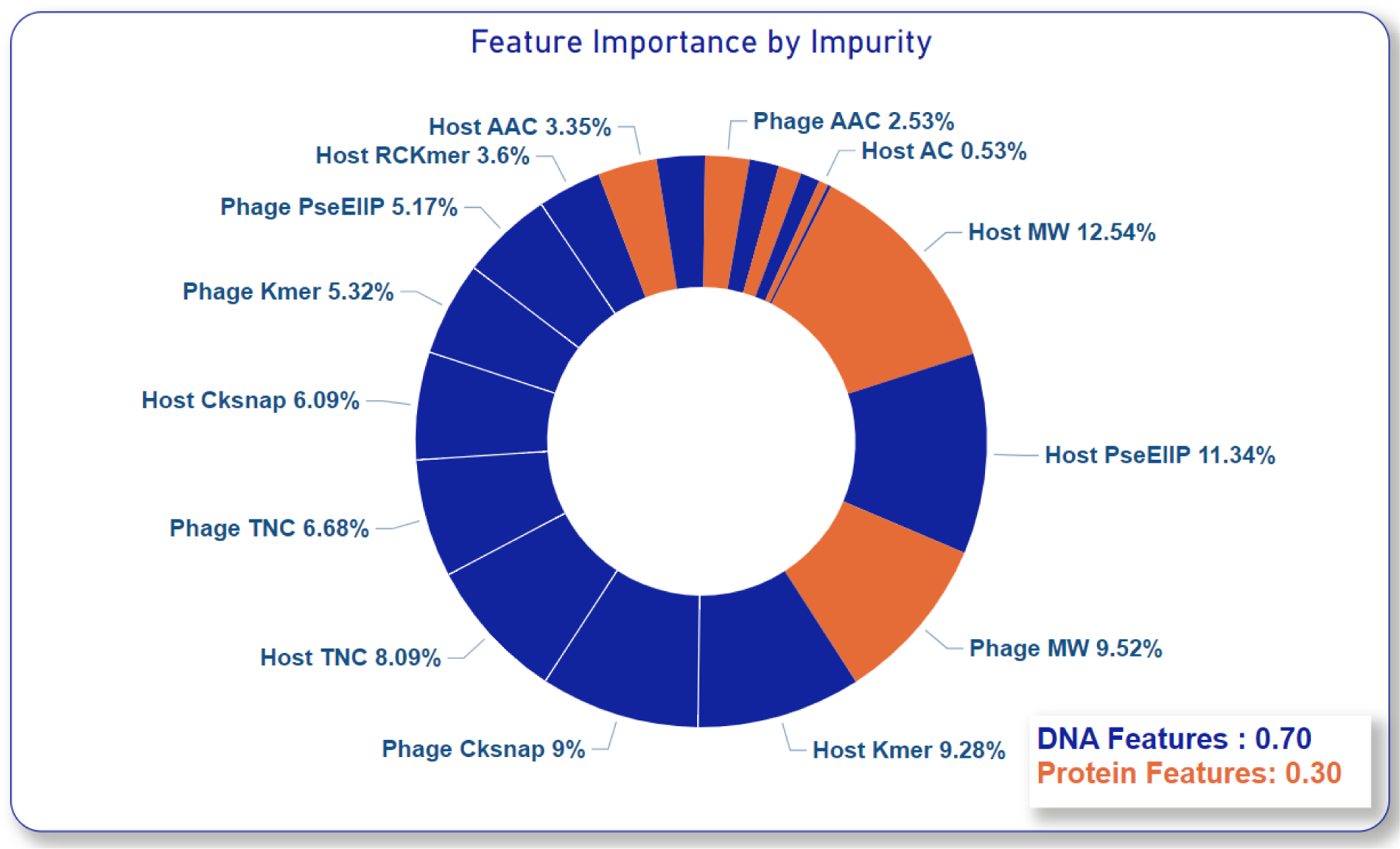
Variable Importance Estimation Using Impurity Method

To assess the prediction sensitivity of the model concerning different groups of hosts within a taxonomic group, further testing was conducted. This involved the following procedures:

1. Exclusion of prevalent ESKAPEE host families from the training set and subsequent testing of prediction accuracies.
2. Inclusion of only prevalent ESKAPEE host families from the training set and subsequent testing of prediction accuracies.

Results from both tests revealed no drastic changes in the prediction accuracies of the composite RF model. This suggests that the model retains its predictive capability and can be effectively employed to predict phages for currently prevalent pathogens. It also suggests that the model retains its performance for hosts that are not common in the current dataset proving that the model will reliably predict phages for any novel hosts that are discovered.

### Improvement

One significant benefit of the composite RF model lies in its versatility, allowing for straightforward expansion to include additional meaningful features that can enhance our understanding of phage–host interactions in future studies. All features following the phage infection cycle such as <u>CRISPR</u> spacers and auxiliary metabolic genes or tRNAs can be included in the feature vectors. This paper only includes feature encodings that are not dependent on the sequence length. Further studies could expand this model to not be restricted by the sequence length.

## Conclusion

Phage therapy is already being used in the personalized treatment of patients for whom traditional antibiotics have failed to work (Yang et al., 2023). Culture-based methods of identifying the host range of a phage are time and labor-intensive and hence can be a bottleneck in the widespread use of phage therapy in modern medicine, especially with the exponential increase in phage classifications by next-gen sequencing methodologies (Klumpp et al., 2012). Recent advancements in computational and bioinformatics tools have made it possible to predict a putative host for a phage with high accuracy, thus reducing the time and effort required to experimentally test a phage’s host range. This paper introduces a novel composite machine learning model that leverages alignment-free methods, by incorporating multiple feature encodings from both nucleotide and protein sequences of phages and hosts and combines it with alignment-based features of alignment scores between phage-phage, phage-host, and host-host. The composite model is not only robust as proven by the 5-fold validation and 70-30 testing but is also interpretable as proven by the ablation studies and variable importance analysis (Li & Zhang, 2022). By incorporating alignment-based scores alongside multiple features from phage and host, the model achieves a notable 5-6% increase in accuracy and AUC. Ablation analysis and variable importance analysis illustrate that nucleotide features contribute more to the performance than proteins, host nucleotides, and proteins have a greater influence than that of phages, and all alignment scores have an equal influence on the performance gain. These results indicate that the composite machine learning model is a promising solution in predicting phage-host interaction. This model can also be used for other classification problems involving nucleotide and protein sequences.

### Data/Code Availability

All data and code used and developed can be found at https://github.com/shrey/CoMPHi.

